# Examination of germline and somatic intercellular bridges in *Hydra vulgaris* reveals insights into the evolutionarily conserved mechanism of intercellular bridge formation

**DOI:** 10.1101/2025.02.19.639158

**Authors:** Kari L. Price, Dyuthi M. Tharakan, Willi Salvenmoser, Kathleen Ayers, Jasmine Mah, Casey Dunn, Bert Hobmayer, Lynn Cooley

## Abstract

Incomplete cytokinesis results in the formation of stable intercellular bridges that have been extensively studied in bilaterians, where they play essential roles in cell-cell communication and coordination of differentiation. However, little is known about their structure and molecular composition in non-bilaterian animals. This study characterizes germline and somatic intercellular bridges in the cnidarian *Hydra vulgaris*, providing insights into their evolutionary origins and functional significance. We identified key conserved components, including KIF23, F-actin, and phosphotyrosine. Notably, we observed microtubule localization within *Hydra* ring canals, suggesting previously unrecognized functions for this cytoskeletal component in intercellular bridge formation. Bioinformatic analyses confirmed the conserved expression of *Kif23* and suggested its role as a molecular marker for identifying ring canal-associated components. EdU incorporation during DNA replication demonstrated that cells connected by ring canals exhibit synchronized cell cycles, which may be critical for the coordination of division and differentiation. Our findings reveal that the molecular and structural features of intercellular bridges in *Hydra* are conserved across evolutionary lineages, highlighting their ancient origins and functional significance in cellular connectivity. The presence of synchronized cell cycles in ring canal-connected cells underscores their role in promoting coordinated cellular behaviors, processes fundamental to multicellular organization. This study provides new perspectives on the evolution of incomplete cytokinesis and establishes a framework for comparative investigations into the diversity and conservation of intercellular bridge mechanisms across metazoans.

## Introduction

Throughout evolution, modifications to the fundamental process of cell division have led to remarkable complexities in cellular architecture and organismal development. In most animal cells, the cell division cycle is punctuated by cytokinesis, ensuring the faithful separation of genetic material, proteins, and organelles into distinct daughter cells. One significant modification to this program is the failure to complete the terminal step of cytokinesis, a process termed abscission, which physically separates nascent cells. Instead, sibling cells remain connected by stable cytoplasmic intercellular bridges, often forming organized syncytial structures known as cysts. Intercellular bridges are widespread across biology where they connect developing animal germ cells and some somatic cell populations (Chaigne and Brunet, 2022). The prevalence of intercellular bridges underscores their evolutionary significance, suggesting they are integral components of cellular communication and coordination essential for multicellular life. However, key questions remain regarding whether the mechanisms driving the formation and maintenance of intercellular bridges are conserved across species, and how these structures originated and evolved to support the complexities of multicellular life.

Cytoplasmic intercellular bridges were first observed in histological sections of interconnected germ cells in testes from the aquatic salamander *Amphiuma* (McGregor, 1899). Subsequent studies in both vertebrate and invertebrate species, including the metazoan *Hydra oligactis*, revealed that intercellular bridges are nearly ubiquitous among germ cells during gamete development, known to connect multipotent stem cells in non-bilaterians, and often link clusters of somatic cells undergoing synchronous differentiation (Burgos and Fawcett, 1955; David, 2012; Dym and Fawcett, 1971; Fawcett et al., 1959; Hobmayer et al., 2012; McLean and Cooley, 2013). Electron microscopy has revealed that both germline and somatic intercellular bridges are short, micrometer-scale channels defined by an electron dense limiting plasma membrane (Burgos and Fawcett, 1955; Koch and King, 1966; Koch and King, 1969). Because of the distinctive ring-shaped structure, these bridges are often referred to as ‘ring canals’, the nomenclature we will use hereafter. Ring canals are crucial for fertility, enabling direct cell-cell communication among germ cells (Chaigne and Brunet, 2022; Gerhold et al., 2022; Greenbaum et al., 2011; Haglund et al., 2011; Lu et al., 2017).

The molecular composition of ring canals and underlying mechanisms of incomplete cytokinesis have primarily been studied in bilaterians, with extensive analyses in the germlines of *Drosophila* and mice, and only recently explored in a non-bilaterian (Greenbaum et al., 2011; Haglund et al., 2011; Price et al., 2023). While there are some compositional differences between species, all bridges examined to date contain the cytokinesis regulator and kinesin-6 family member, KIF23. As a subunit of the centralspindlin complex, KIF23 plays a crucial role throughout all stages of cytokinesis, linking the plane of cell division with the anaphase spindle, driving the formation of the central spindle, and serving as a major component of the midbody - a structure critical for abscission of daughter cells (Glotzer, 2009; White and Glotzer, 2012). During germ cell incomplete cytokinesis in *Drosophila*, mice, and the cnidarian *Hydra vulgaris*, KIF23 and its associated proteins organize the formation of a midbody, but rather than initiating abscission, the KIF23-rich midbody is remodeled to form a ring canal with an open lumen (Price et al., 2023). Our observations of midbody reorganization in distantly related animals suggests that the cellular and molecular mechanisms of incomplete cytokinesis and ring canal formation may be conserved among metazoans.

The presence of ring canals in both germ and somatic cell populations points to the broad functional significance of these structures in diverse cell types and developmental programs. Furthermore, germ cells and multipotent stem cells in non-bilaterian animals like *Hydra* share key molecular characteristics, including the expression of core germ cell genes like Nanos, Vasa, and Piwi (Alié et al., 2011; Hemmrich et al., 2012; Juliano et al., 2011; Juliano et al., 2014; Spradling, 2024). These shared features suggest that ring canal formation first emerged in ancestral multipotent stem cell populations and directly contributed to the evolution of multicellularity and germline development by facilitating cooperative functions within a shared environment (Spradling, 2024). However, no studies have directly examined ring canals in these different cellular contexts in an early animal.

The deep phylogenetic relationship between cnidarians and bilaterians offers a unique opportunity to explore the evolution of ring canal formation and function. In this study, we integrated what is known about ring canals in bilaterian animals with new knowledge of ring canal composition in *Hydra vulgaris* to identify shared molecular components. This approach allowed us to uncover common features that likely existed before the divergence of cnidarians and bilaterians. We characterized *Hydra* germ cell and somatic ring canals with immunofluorescence and electron microscopy and found that all cells in the interstitial stem cell lineage are connected by ring canals that are invariant in size at just under one micron in diameter. Furthermore, we found that all ring canal-connected cells display cell cycle synchronicity and coordinated differentiation. We analyzed published single cell-RNA sequencing datasets to bioinformatically infer candidate ring canal components and conserved interactions. Interestingly, we find that enrichment of *kif23* transcripts accurately predicts the cells linked by KIF23-labeled ring canals in *Hydra* and those suspected to be linked in different taxa. These data suggest that KIF23 is part of the core suite of components required for ring canal formation in the earliest animals and raise important questions about how *kif23* expression is differentially regulated in cells that undergo incomplete cytokinesis.

## Results

### Examining ring canals in the cnidarian Hydra vulgaris

Given the critical role of KIF23 in cytokinesis and its presence in ring canals in various species, we sought to investigate the extent of its conservation across evolutionarily distant lineages and gain insight into the earliest ring canals. For this, we turned to the simple cnidarian *Hydra vulgaris*. In addition to its phylogenetic position, *Hydra* contains well-studied multipotent stem cells, called interstitial stem cells (ISCs), which are often observed as pairs linked by cytoplasmic bridges (David, 1973; Hobmayer et al., 2012; Holstein and David, 1990). In contrast to the unipotent ectodermal and endodermal epithelial stem cells that give rise to the two layers of epithelium (Figure 1A), *Hydra* ISCs have the capacity to produce all non-epithelial cell types required for tissue homeostasis and sexual reproduction including both gametes and the somatic cell types of neurons, gland cells, and nematocytes (Figure 1B). For this study, we focused on two ISC-derived lineages, the nematocytes and male gametes, as these cell types have long been understood to form interconnected nests of cells. ISCs can divide either symmetrically to produce a pair of linked ISCs or asymmetrically to produce a single founder nematoblast, the precursor to the nematocyte and a major cell type used both to capture food and in defense. The nematoblast and male and female germ cells undergo successive rounds of incomplete cytokinesis, resulting in interconnected nests of 4, 8, 16 and 32 cells in the nematocyte lineage and clusters of germ cells in the germline (David, 1973; David, 2012; Hobmayer et al., 2012; Holstein and David, 1990).

**Figure 1.**
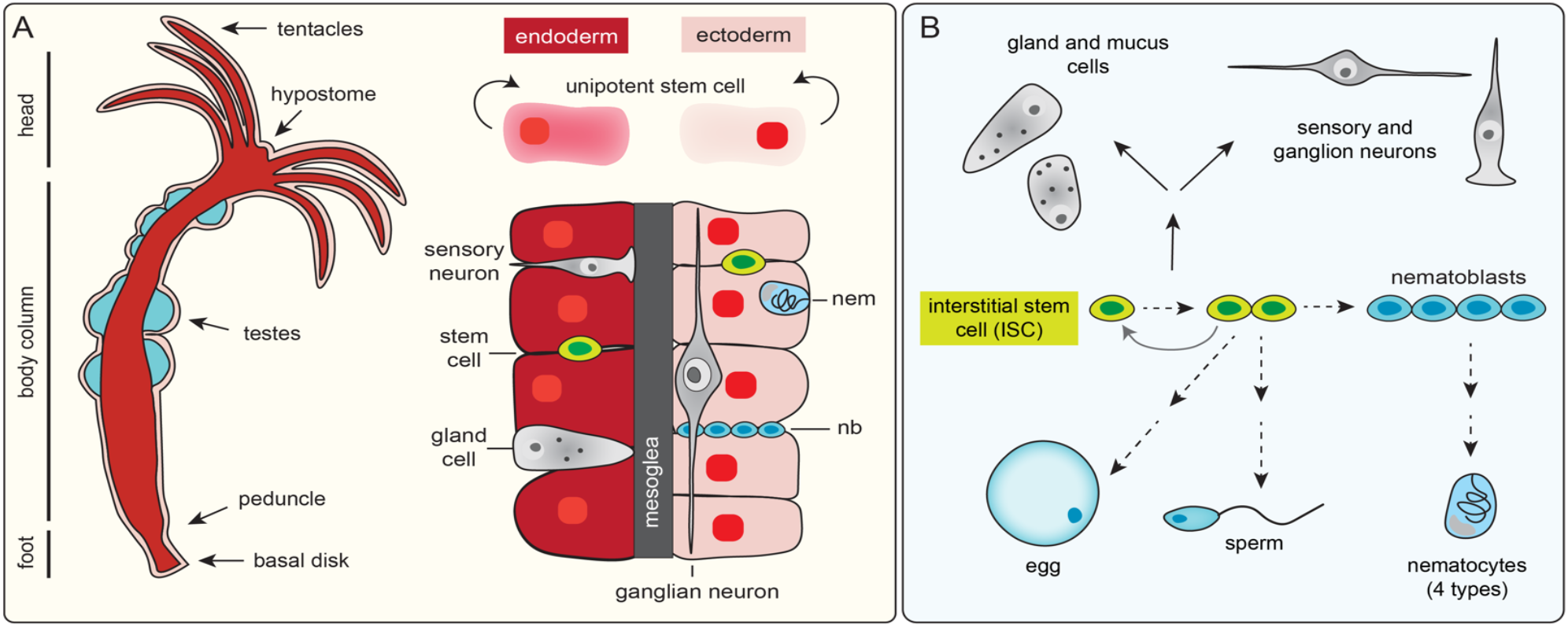
Schematic of the cell types and cellular differentiation pathways in *Hydra vulgaris*. (A) The *Hydra* polyp has a tubular, radially symmetrical body with an adhesive foot at the aboral end (basal disk) for attachment to substrates and a mouth at the opposite end that is surrounded by tentacles. The mouth is positioned at the tip of the cone-shaped hypostome. The body column is comprised of two layers of epithelial cells - the endoderm (red) and ectoderm (pink) - two concentric cell layers that are separated by an extracellular matrix called the mesoglea. The endoderm and ectoderm have stem cell properties and can produce additional ectodermal or endodermal epithelial cells that are displaced be cell divisions along the oral-aboral axis to occupy the hypostome, tentacles, and foot. The testes are situated in the ectodermal epithelium. At the interstices of the ectodermal epithelial cells are the interstitial stem cells (ISC, ‘stem cell’, cyan) that give rise to differentiated daughter cells (shown in cyan and grey). Nematoblast, nb; nematocyte, nem. (B) ISCs are multipotent stem cells, often appearing in pairs, whose products yield non-epithelial cell types in the animal. Proliferating ISCs have the potential to divide either symmetrically or asymmetrically to produce both terminally differentiated somatic cells and gametes. In nematocytes and the male and female germlines, precursor cells divide incompletely, remaining linked by intercellular bridges during differentiation (as indicated by the dotted arrows).

The abundance of ring canals in the interstitial cell lineages in *Hydra* provided an opportunity to explore the composition and function of both somatic and germline ring canals and a window into the ancestral characteristics of these structures. In a previous study, we generated and validated a polyclonal antibody that recognizes the *Hydra* homolog of KIF23 (HyKIF23; (Price et al., 2023)) and focused on the localization of HyKIF23 in the male germline. Here we have expanded our focus to include the somatic cells connected by ring canals. In intact polyps, we found HyKIF23-labeled ring canals connecting pairs of ISCs, nests of nematoblasts, and immature nematocytes along the length of the body axis (Figure 2A-D). We found no evidence of ring canal-connected syncytia in the tentacles, where fully differentiated and individualized nematocytes reside (Figure 2E). To confirm the identity of each cell type, we imaged macerated tissues (Figure 2F-I) or identified cells by the presence or absence of the stem cell markers Hywi (*Hydra* Piwi; Juliano et al., 2014) or GFP under the control of Cnnos (*Hydra* Nanos; Hemmrich et al., 2012) in intact animals (Supplemental Figure 1). In contrast to individual ISCs where ring canals were never observed, every ISC pair identified was connected by a HyKIF23-labeled ring canal (Figure 2G; n=19). Similarly, we observed ring canals at all stages of nematoblast differentiation and in all terminally divided, but not fully differentiated nematocytes (Figure 2C-E, 2H-I’). As expected, neither epithelial (Supplementary Figure 2), nerve, nor gland cells were connected by ring canals.

**Figure 2.**
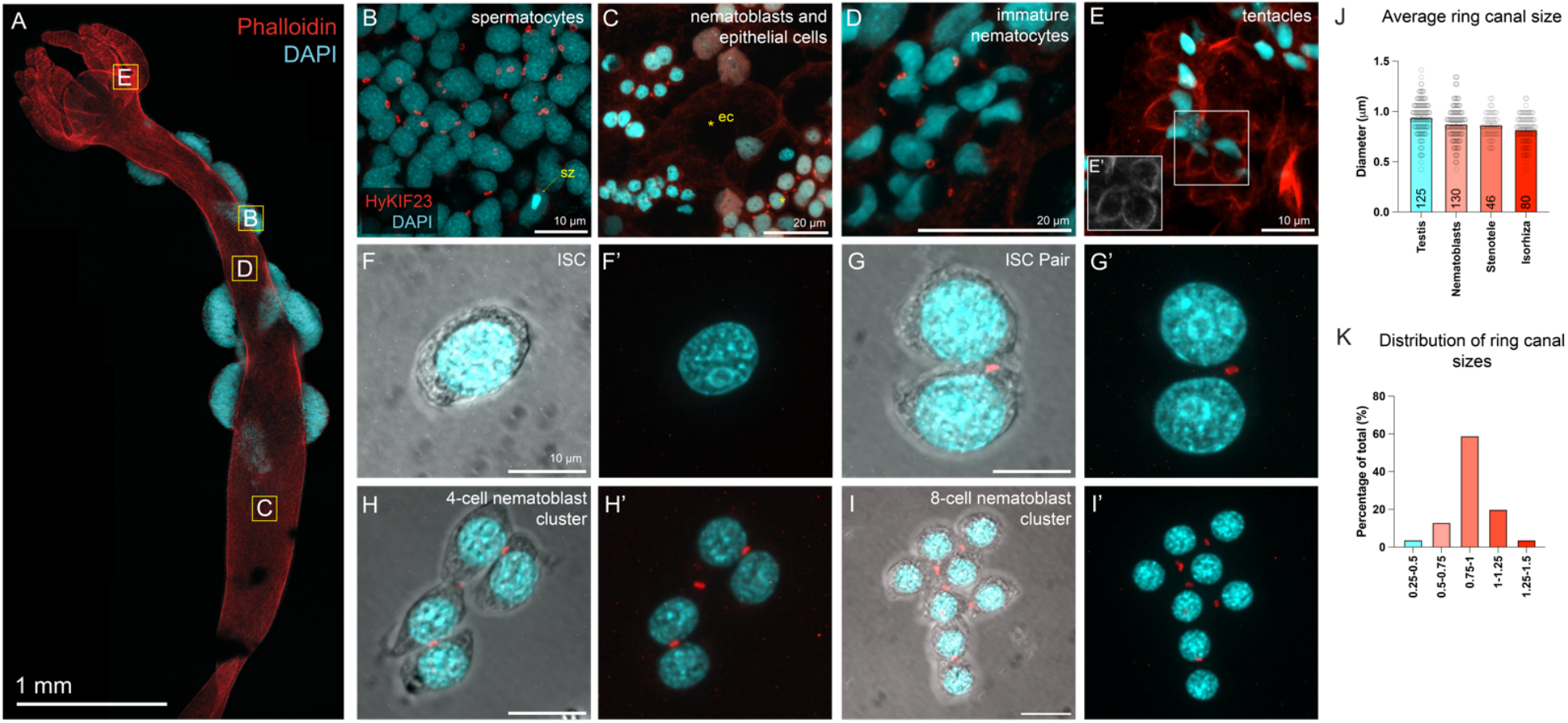
HyKIF23-labeled ring canals connect a subset of cells of the interstitial lineage. (A) A male *Hydra* polyp. Regions in high magnification images in B-E: B, testis; C, D, body column; E, tentacles. Phalloidin, red; nuclei, cyan. (B-E) Localization of HyKIF23 to ring canals in the following cell types: (B) Primary spermatocytes in the *Hydra* testis. A haploid spermatozoon is visible in the field (‘sz’, yellow arrow). (C) A field of epithelial cells and nematoblasts; HyKIF23-labeled ring canals are only present between clusters of nematoblasts and not between the larger epithelial cells (‘ec’, yellow asterisks). (D-) HyKIF23 localizes to ring canals that connect the immature nematocytes (D), but do not connect mature nematocytes in the tentacles (E). (F-I’) Macerated cells with intercellular bridges. (J) Average ring canal diameters in germline and somatic cells (p=0.0025, Kruskal-Wallis test). (K) Distribution of germline and somatic ring canal diameters measured in syncytial clusters from macerated tissue preparations.

Having established that HyKIF23-labeled ring canals are present in multiple somatic cell types, we measured ring canal diameters across the diverse interstitial cell lineage to determine any difference by cell type. While the difference between all cell types was statistically significant (p=0.0025, Kruskal-Wallis test), the observed differences in means is smaller than the resolution limit of light microscopy, suggesting that the variability in ring canal diameter between germ and somatic cells is not biologically significant (Figure 2J). This is in stark contrast to observations in *Drosophila* where ring canals vary in size up to 40-fold depending on the cell type, reflecting their functional differences (Airoldi et al., 2011; Kramerova and Kramerov, 1999; McLean and Cooley, 2013; Poodry and Schneiderman, 1970). Our data suggest that ring canals in *Hydra* may be functionally similar, if not identical, across all connected cell types.

Whereas ring canal connections between ISC pairs have been described using brightfield illumination of macerated cells, to our knowledge no studies have examined these structures with electron microscopy. We therefore examined ring canals in pairs of ISCs with transmission electron microscopy and compared this ultrastructure with those of ring canals in the male germline and in immature nematocytes (Figure 3). Ring canals that connect ISC pairs are marked by an electron dense limiting membrane that is continuous with the plasma membrane of the connected sibling cells (Figure 3A). All ring canals observed in our electron micrographs formed a tire rim-shape similar to descriptions of ovarian ring canals in *Drosophila* (Robinson et al., 1994). By comparing ring canals in both germ and somatic cell types, our data reveal that the size, shape, and ultrastructure of ring canals is highly conserved across these different lineages. Together, these findings suggest that despite the functional diversity of cell types in *Hydra*, the structural properties of ring canals, including size, shape, and ultrastructure, are highly conserved across both somatic and germline lineages, reflecting a shared mechanism of intercellular communication and coordination.

**Figure 3.**
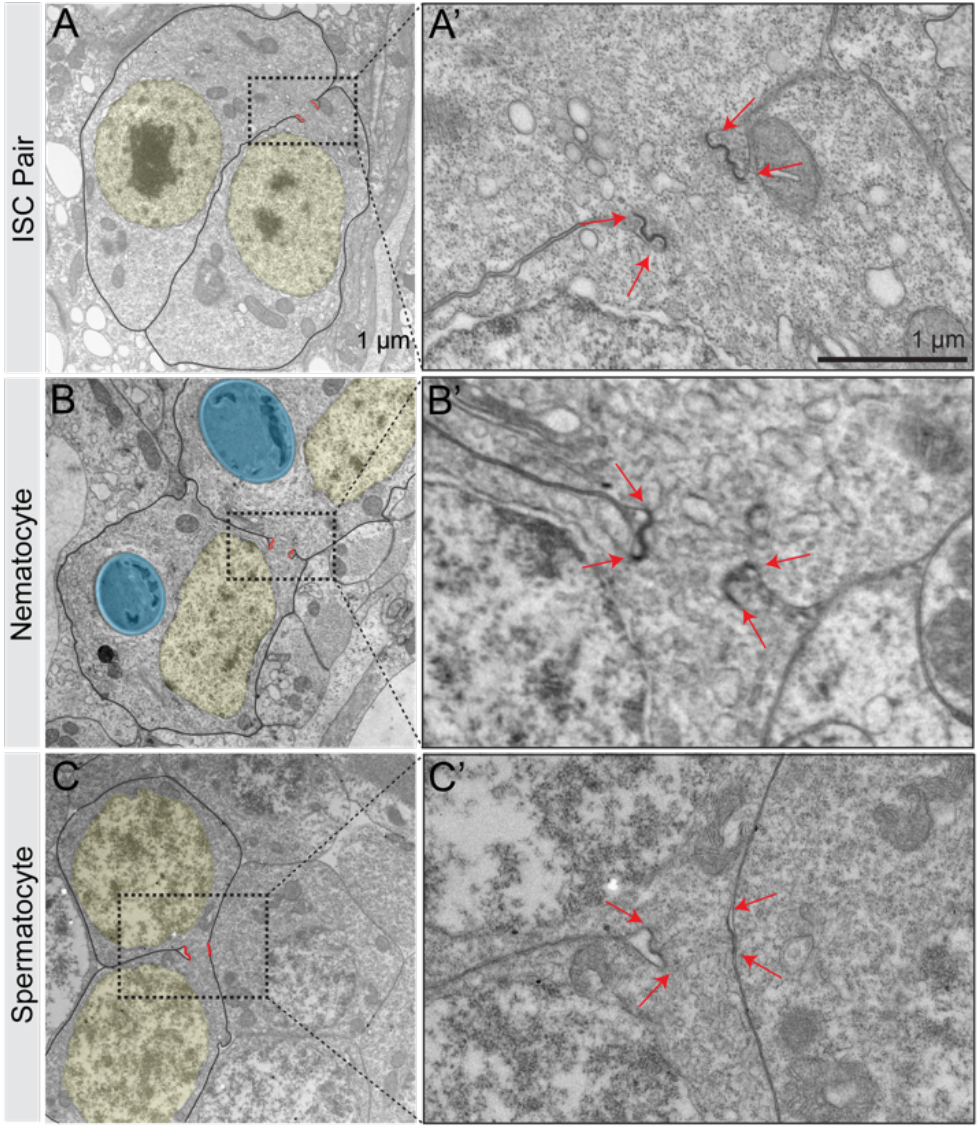
Ultrastructure of a *Hydra* interstitial stem cell (ISC) pair (A-A’), differentiating nematocytes in a nest (B-B’), and primary spermatocytes (C-C’). Nuclei, yellow; ring canals, red; nematocyst capsule, blue. The boundaries of the electron dense-limiting membrane of the ring canal are marked by red arrows.

### Cell cycles are synchronous among linked cells

Early investigations noted the presence of ring canals connecting groups of cells undergoing synchronous differentiation (Fawcett et al., 1959). Subsequent studies in *Drosophila* and rodent models revealed that ring canals facilitate the sharing of cytoplasmic components and equilibration of protein between transcriptionally mosaic cells, likely promoting coordinated differentiation and cell division programs as well as buffering inconsistent gene expression across syncytial clusters (McLean and Cooley, 2013; Ventelä et al., 2003).

To test the hypothesis that ring canals coordinate cell cycle synchronization in a non-bilaterian, we performed EdU incorporation assays to evaluate cell cycle dynamics in the syncytial cell types in *Hydra vulgaris*. We soaked adult polyps continuously for 48 hours in 100 μM EdU and then detected proliferation patterns on whole mounts stained with anti-HyKIF23 to visualize ring canals. Consistent with previous reports, we observed few cycling cells in the hypostome and tentacles (Buzgariu et al., 2014) (Figure 4A). In contrast, and as expected, we observed high numbers of EdU-positive nuclei spanning the length of the body column with varying degrees of EdU incorporation (Figure 4B-F). In all cases, EdU-positive nuclei that were within five microns of another EdU-positive nucleus were linked by a HyKIF23-labeled ring canal and displayed identical patterns of EdU-incorporation. These results suggest that ring canals in *Hydra* play a critical role in coordinating cell cycle dynamics, ensuring synchronized proliferation within syncytial clusters, a function that may be conserved across both bilaterian and non-bilaterian lineages.

**Figure 4.**
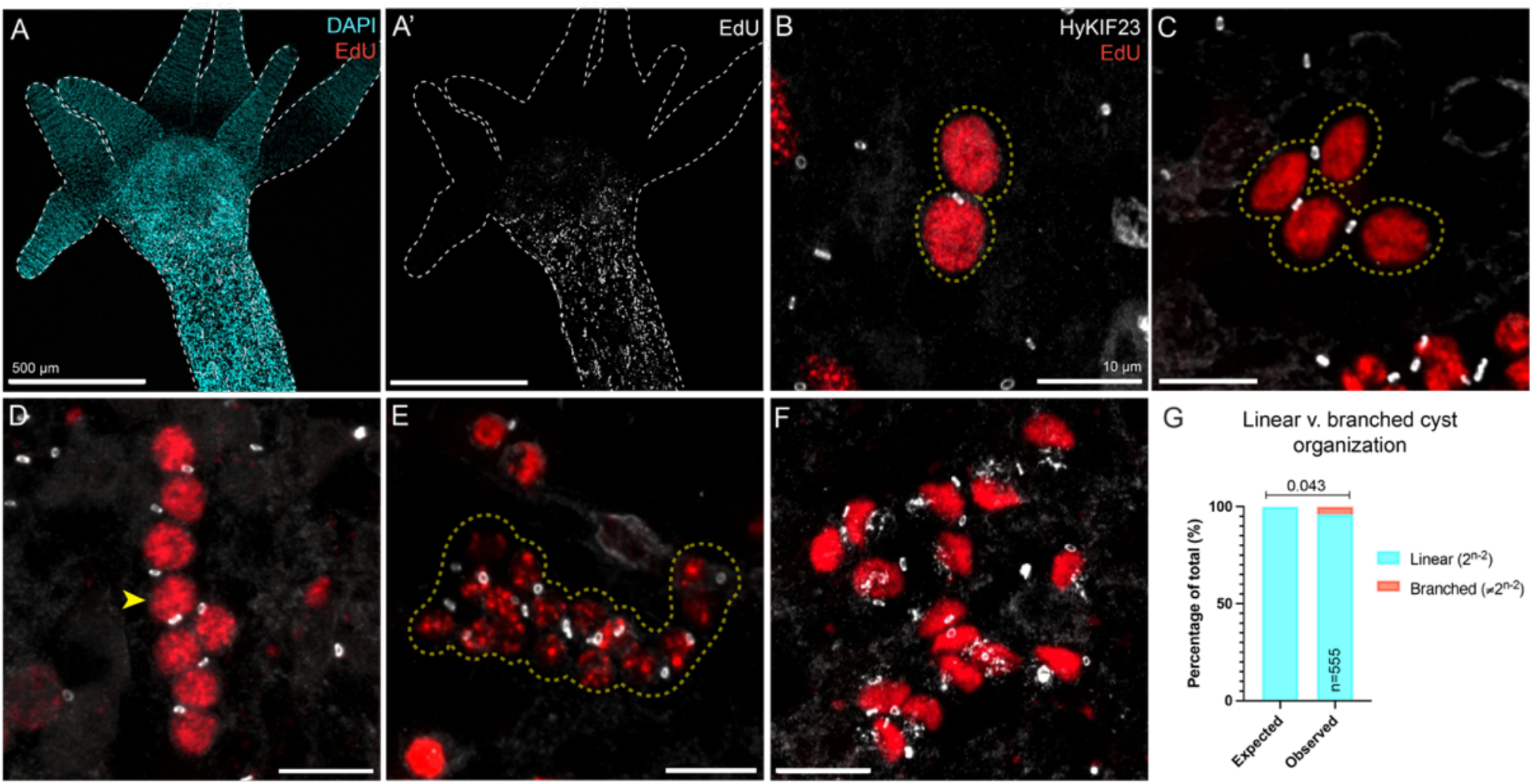
The cell cycles of cells connected by ring canals are highly synchronized. (A-A’) *Hydra* polyp with EdU- and DAPI-labeled nuclei. (B-F) EdU-positive nuclei and immunostaining of anti-HyKIF23 in intact animals. (B) A pair of interstitial stem cells. (C) A four-cell nest of nematoblasts. (D) An eight-cell nest of nematoblasts, with a nucleus that is connected to three neighboring cells by three ring canals (yellow arrowhead). (E) A 16-cell nest of nematoblasts. (F) A 16-cell nest of differentiated nematocytes. (G) Quantification of nuclei-ring canal connections in 8-cell nests.

The mosaic EdU labeling also offered an opportunity to evaluate nest architecture and connectivity. Drawings of nematoblast nests have suggested they are organized in linear arrays with most cells having two ring canals, but the exact branch patterns have not been reported (Spradling, 2024). We therefore quantified the number of nests that contained cells with more than two ring canal connections. If arranged in a linear fashion, the number of ring canals in a single nest would be equal to 2^n-2^; deviations from this number would represent a branched pattern of connectivity. As branching would only be evident in nests of eight cells or more, we restricted our quantification to eight-cell nematoblast nests (Figure 4D and G). In eight-cell nematoblast nests, 96% of syncytia were arranged in a linear pattern with each nucleus connected to no more than two neighboring nuclei. The remaining 4% of syncytia often contained a single nucleus with three ring canals, resulting in a minor branched architecture (Figure 4D, yellow arrowhead). These data indicate that nests in *Hydra* predominantly follow a linear organizational pattern, closely resembling the linear organization of mouse spermatogonia and spermatocytes (Yoshida, 2016).

### Ring canal protein composition resembles that of more recently evolved eukaryotes, with exception

Informed by immunofluorescent analyses of ring canal composition in *Drosophila* and mouse, we sought to identify additional ring canal components in *Hydra*. As there are few *Hydra-*specific antibodies, we examined the localization of proteins for which there are commercially available, cross-reactive antibodies and reagents. F-actin is a component of female ring canals in *Drosophila*, where it plays a crucial role in maintaining their structure and function in the ovary, and is also found in male intercellular bridges in mice, likely contributing to their stability and synchronization of germ cell development (Cooley and Theurkauf, 1994; Greenbaum et al., 2007). In *Drosophila* male germline ring canals, F-actin is a less-prominent component and they instead contain a septin cytoskeleton (Hime et al., 1996). To determine whether F-actin was a component of *Hydra* germline and somatic ring canals, we fixed and co-stained adult polyps with anti-HyKIF23 and Phalloidin. In all germline and nematoblast cells, F-actin co-localized with HyKIF23 in the mature ring canal (Figure 5A-D). Line scans of F-actin fluorescence intensities relative to HyKIF23 revealed extensive colocalization in the ring canal, with an accumulation of F-actin at the inner diameter of the ring canal (Figure 5G), similar to ring canals in the *Drosophila* female germline. These findings reveal that the spatial organization of F-actin and HyKIF23 in Hydra ring canals mirrors that of *Drosophila*, suggesting a conserved structural role for F-actin in ring canal stability and function across evolution.

**Figure 5.**
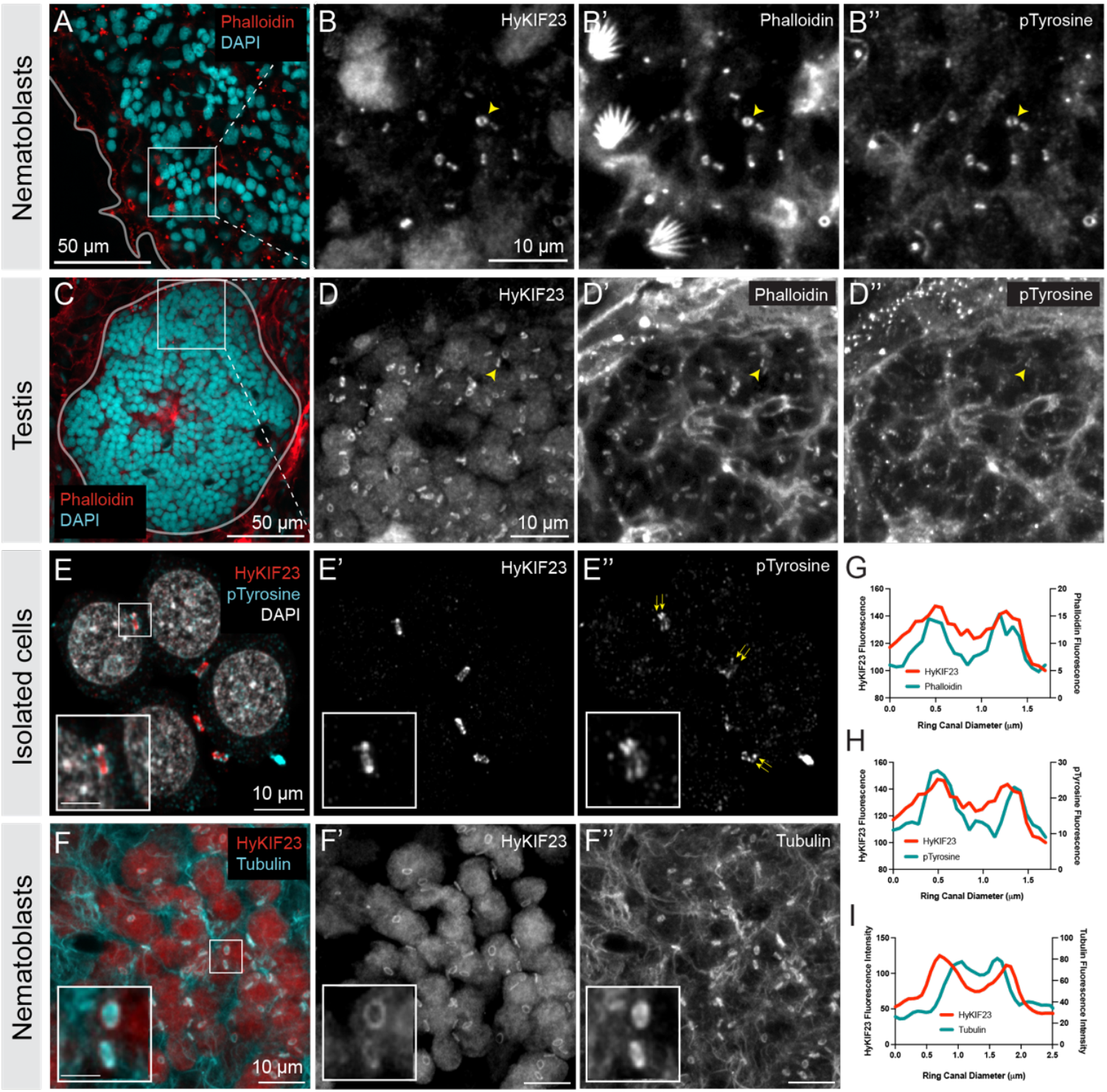
*Hydra vulgaris* molecular ring canal composition. (A-B’’) A cluster of nematoblasts immunolabeled with anti-HyKIF23 to label ring canals, Phalloidin to visualize F-actin, and anti-pTyrosine to mark the pTyrosine epitopes in the ring canal. The same ring canal is marked with a yellow arrowhead. (C-D’’) A cluster of spermatocytes in the *Hydra* testis labeled with the same antibodies as in A-B’’. The yellow arrowhead marks a ring that is triple positive for HyKIF23, Phalloidin, and pTyrosine. (E-E’’) Syncytial cells from macerated preparations to better visualize pTyrosine localization. Insets reveal that Phosphotyrosine signal is present as two rings that form on either side of the central HyKIF23-labeled ring canal, similar to previous descriptions of Phosphotyrosine labeling in Drosophila. (F) Ring canals connecting nematoblasts are positive for Alpha-tubulin signal. (G-I) Line scans of representative ring canals labeled with phalloidin (G), pTyrosine (H), and tubulin (I).

In *Drosophila*, phosphotyrosine epitopes are a well-characterized presence in germline and somatic ring canals and one of few components that have not been shown to be directly required for cytokinesis. Examination of phosphotyrosine in *Hydra* ring canals revealed a positive signal in both germline and somatic ring canals (Figure 5). We observed phosphotyrosine on all cell membranes with marked enrichment on ring canals that were co-stained for HyKIF23. Upon closer examination, we found that phosphotyrosine did not co-localize with HyKIF23 in the ring canal, but rather localized to ring-shaped structures on both sides of the central HyKIF23 signal (Figure 5E). This staining pattern is consistent with the staining patterns observed in *Drosophila* male germline cells (Eikenes et al., 2015), but which have not been described in the mouse male germline. These findings suggest that the localization of phosphotyrosine in *Hydra* ring canals mirrors patterns observed in *Drosophila* male and female germlines, providing further evidence of conserved roles for phosphotyrosine across species and cell types in the structure and function of intercellular bridges.

In addition to investigating well-established components like F-actin and phosphotyrosine, we examined microtubules. Unlike F-actin and phosphotyrosine, there is no precedent for microtubule localization to ring canals in other species. However, the localization of KIF23, a known microtubule motor, to all ring canals begs the question of whether microtubules may have been involved in the structure or function of these intercellular bridges during evolution. We fixed and stained *Hydra* polyps with an anti-alpha tubulin antibody. Although we also examined the testis, the abundance of tubulin signal in this tissue made it challenging to resolve ring canals from the dense and overlapping sperm axonemes. Therefore, we focused on the nematoblast lineages as their linear arrangement along the body axis simplified imaging and allowed for clearer visualization of ring canal components. We found a positive alpha-tubulin signal on ring canals in a subset of cysts (∼13%) within the nematoblast lineage (Figure 5F-F”). In those tubulin-positive ring canals, the signal was always on the luminal side of the ring canal and was present as either lining the ring canal or with a more diffuse disc-shaped localization filling the ring canal lumen (Figure 5I). Interestingly, the diffuse tubulin signal that we observed was distinct from that of mitotic spindle remnants, which form a compact bundle that we have shown to occupy the nascent ring canal lumen in *Drosophila* (Price et al., 2023). The tubulin-containing ring canals in *Hydra* are likely nascent ring canals, with the retention of microtubules within the lumen representing a transient and newly observed stage of ring canal biogenesis in cnidarians. This observation supports the idea that microtubules might play a previously unrecognized role in ring canal architecture in early-diverging metazoans, potentially pointing to an ancient origin of this cytoskeletal component in intercellular bridge formation.

### HyKif23 gene expression is enriched in cell types connected by ring canals

To complement our cell biological study of *Hydra* ring canal proteins, we analyzed publicly available single-cell RNA sequencing datasets in *Hydra* (Cazet et al., 2023; Siebert et al., 2019) to examine *hyKif23* gene expression patterns. Consistent with our results from fixed immunofluorescence experiments, *hyKif23* (annotated as G003313) gene expression was highest in cells of the interstitial cell lineage, including the interstitial stem cells, the male and female germ cells, early nematoblasts, and neural progenitor cell populations (Figure 6A, B). As expected, this expression pattern was mirrored by the *Hydra* homolog of RacGAP (G020158), a known and conserved binding partner of KIF23 and the only other subunit in the centralspindlin complex. Given the established role of the centralspindlin complex in mitotic cytokinesis, the high levels of transcripts observed in these cell populations could be attributed to the active involvement of *hyKif23* and *hyRacGAP* in cell division. When we compared the gene expression profiles of *hyKif23* to that of the key mitotic cell cycle regulator *Cyclin B* (G001809), we observed extensive overlap as would be expected of two genes involved in mitotic cell cycle progression (Figure 6D). However, while all cell types that expressed *hyKif23* also expressed *Cyclin B*, these two genes were not equally expressed across all cell lineages; for example, nematoblast cells undergoing incomplete cytokinesis to produce ring canals expressed higher levels of *hyKif23* and *Cyclin B* than cells that complete cytokinesis (e.g. ectodermal and endodermal cells) implying that enrichment of *hyKif23* is not strictly linked to its mitotic activity.

**Figure 6.**
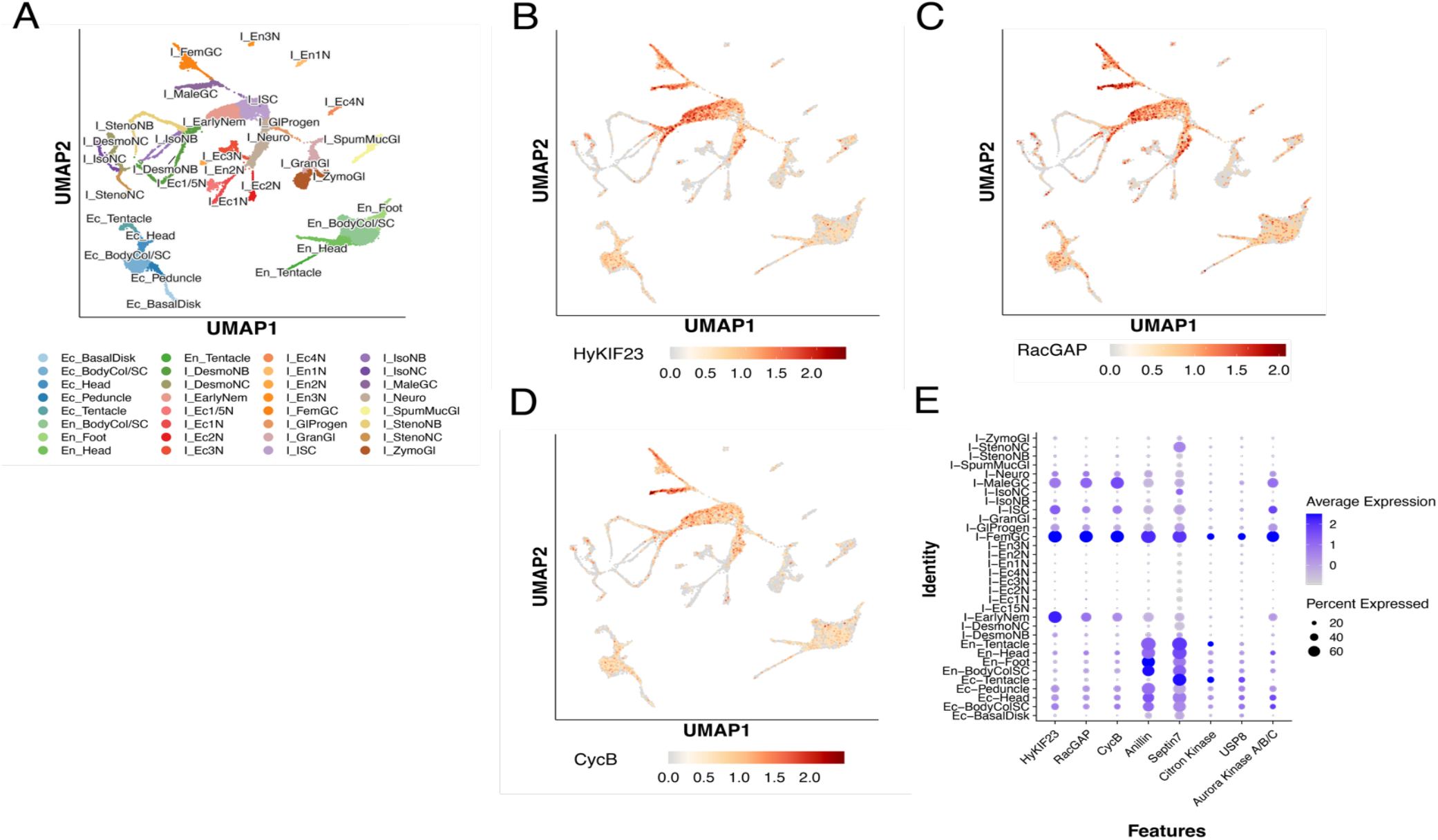
(A) UMAP of clustered cell types from Cazet et al. (2023). (B-D) Expression profiles of *HyKif23, hyRacGAP*, and *hyCyclin B*. (D) Dot plot of *HyKif23, hyRacGAP*, and *hyCyclin B* and potential ring canal homologs identified through BLAST of known ring canal genes in *Drosophila*.

The agreement between *hyKif23* gene expression and HyKIF23 protein localization suggest that enrichment of *hyKif23* may not only serve as a robust predictor of cell types linked by ring canals but also offers an opportunity for the bioinformatic inference of other potential ring canal components. By comparing these profiles, we aimed to identify additional genes in *Hydra* that might play a role in ring canal development, thereby expanding our understanding of the molecular machinery underlying this conserved process across diverse organisms. We analyzed the same single-cell RNA sequencing datasets, querying the expression profiles of *Hydra* homologs of genes previously identified in *Drosophila* and mouse to encode for ring canal components. We performed a BLAST search to first identify *Hydra* homologs of known ring canal resident or promoting components, including Anillin, Septin 7 (*Drosophila* Peanut), Citron Kinase (*Drosophila* Sticky), USP8, and Aurora Kinase A/B/C (Figure 6E). The homologs of these genes are expressed in many of the same cell types as *hyKif23*, with similar expression profiles. Interestingly, *Anillin* and *Septin 7* were also enriched in the endodermal and ectodermal cell populations. As these cell types undergo complete cytokinesis, these expression patterns may reflect the role of Anillin and Septin 7 in the coordination of cell division more generally or as a structural component of differentiated cells. Despite this, the enrichment of ring canal homologs in the same *hyKif23-* expressing cells suggests that they may contribute to ring canal biogenesis and may be ring canal candidate proteins in early animal syncytial cells. We conducted the same analysis in other cnidarian species for which single-cell RNA sequencing datasets have been published (Chari et al., 2021; Schnitzler et al., 2024; Steger et al., 2022). We observed that, as in *Hydra* and *Drosophila*, undifferentiated and/or germ cells displayed higher levels of *Kif23* transcripts (Supplemental Figure 3). Together, these data suggest the possibility that enrichment of *Kif23* may be a marker of cell types that undergo incomplete cytokinesis to produce ring canals.

## Discussion

Most studies of ring canals have focused on bilaterians, particularly in the germlines of *Drosophila* and mouse, where the molecular composition and the mechanisms of incomplete cytokinesis and ring canal formation are relatively well-characterized. Here, we expand this field of study to the non-bilaterian and simple cnidarian *Hydra vulgaris*, providing the first detailed examination of ring canal abundance, morphology, and composition in an early-diverging metazoan. Our findings reveal that *Hydra* shares key conserved features of ring canals with bilaterians, while also uncovering novel aspects of ring canal composition and biogenesis that may reflect ancestral mechanisms and inform our understanding of the origins of animal multicellularity.

### New insights into ring canal composition and size

Our findings expand the understanding of ring canal composition by revealing both conserved and unique molecular features. Key components like KIF23 and F-actin, which are conserved in mice and Zebrafish germline ring canals and *Drosophila* germline and somatic ring canals (Bertho et al., 2021; Greenbaum et al., 2011; Haglund et al., 2010; Haglund et al., 2011; Robinson and Cooley, 1996; Russell et al., 1987) play crucial roles in their formation and stability. Here, we show that these same components comprise both germline and somatic ring canals in *Hydra*, indicating that the molecular toolkit for ring canal formation is deeply conserved and may have been established early in evolution to support intercellular connectivity.

The relative sizes and composition of germline and somatic ring canals across species reflect their distinct roles in cellular and developmental processes. Larger germline ring canals may accommodate increased cytoplasmic transfer to support gamete development, while somatic ring canals, often smaller, may facilitate cell-to-cell communication or promote the organization of cells within a tissue. Understanding these size differences may provide insights into how ring canal size and function are tailored to meet the specific demands of germline and somatic cells, offering a window into the evolutionary and functional diversity of these structures. Interestingly, in *Hydra*, the diameters of germline and somatic ring canals are similar, a feature distinct from *Drosophila*, where germline ring canals are many times (4-to 40-fold) larger than their somatic counterparts. This uniformity in Hydra suggests possible differences in the functional requirements or mechanical constraints placed on ring canals early in evolution.

Phosphotyrosine epitopes - a prominent ring canal component in *Drosophila* somatic and germline ring canals (Eikenes et al., 2015; Robinson and Cooley, 1996) - are also present in both somatic and germline ring canals in *Hydra vulgaris*. To date, phosphotyrosine epitopes have not been described in zebrafish or mouse ring canals, suggesting that this feature represents an ancestral characteristic that was lost as more derived lineages evolved alternative mechanisms to effect incomplete cytokinesis. These mechanisms could include changes in cell signaling pathways; differences in the assembly, stabilization, and/or regulation of the cytoskeleton as a result of differences in tyrosine phosphorylation; absence or replacement of specific tyrosine-phosphorylated proteins within ring canals; or even metabolic differences. Proteomic studies of ring canal composition in *Hydra vulgaris* as compared to zebrafish or mice is poised to reveal the molecular substitutes or innovations that fulfill this role in mammalian germlines and shed light on the evolution of incomplete cytokinesis.

In addition to KIF23, F-actin, and phosphotyrosine, our study reveals two distinct populations of microtubules within *Hydra* ring canals. The first population, consistent with observations in bilaterians, consists of compact bundles of spindle-remnant microtubules that traverse the nascent ring canal lumen. These remnants are associated with the KIF23-labeled midbody that undergoes dramatic reorganization to form the nascent ring canal (Price et al., 2023). The second population of microtubules is disc shaped and fills the nascent lumen of the ring canal and has not been described in bilaterians. While the precise timing of the formation of this second population remains unclear without live imaging, the absence of compact spindle microtubules and the presence of the disc-shaped structure within the ring canal suggest that the spindle remnants may be disassembled and reorganized into this distinct arrangement. This population of microtubules also appears to be highly transient, as it is observed with low frequency in fixed samples. The disc-shaped microtubule structure may represent an ancestral configuration of microtubules within ring canals, reflecting an earlier role for stabilizing and organizing the cytoskeleton during incomplete cytokinesis. Furthermore, the microtubule motor activity of KIF23 at these structures may be central to their assembly.

### Differential regulation of KIF23 in cells undergoing incomplete cytokinesis

By integrating analyses of published single-cell RNA sequencing datasets with our cell biological investigation of ring canals, we discovered that HyKIF23 and KIF23 homologs in other species are enriched in cells that form ring canals (Figure 6B, Supplemental Figure 3). Unlike cells that complete cytokinesis where KIF23 expression is lower, the elevated levels in syncytial cells suggest a regulatory mechanism tailored to support intercellular connectivity. In a previous study, our work suggested that reduced KIF23 protein levels in the germ cell midbody impaired ring canal formation (Price et al., 2023) and here we extend these findings by showing that elevated KIF23 mRNA levels correlate with the protein abundance necessary for incomplete cytokinesis. We hypothesized that higher KIF23 expression facilitates the formation of the large midbodies that are necessary for the subsequent remodeling into ring canals (Price et al., 2023). This relationship underscores the importance of precise regulation of KIF23 at both the transcriptional and protein levels in determining cytokinesis outcomes, with increased KIF23 favoring the stabilization and subsequent reorganization of midbody structures into ring canals. These observations suggest that KIF23 not only supports the structural requirements for ring canal formation but may also act as a molecular determinant of the transition between complete and incomplete cytokinesis. Whether mRNA stability or increased transcription underlie the elevated expression of KIF23 in syncytial cells is a subject of future research.

With this in mind, it is then intriguing to consider the outcomes of unintended incomplete cytokinesis events, like those associated with cancer. Elevated levels of KIF23 are frequently observed in the context of human cancer, with overexpression linked to poor prognoses and heightened aggressiveness (The Cancer Genome Atlas, Castoldi et al., 2024; Ji et al., 2021; Kato et al., 2016; Li et al., 2022a; Xu et al., 2023). High levels of KIF23 may be sufficient to disrupt the normal progression of cytokinesis without fully recapitulating the formation of *bona fide* ring canals, resulting in abnormal intercellular bridges or incomplete divisions that ultimately contribute to tumor progression and resistance to therapies. Whether such bridges exist in the context of cancer remains to be seen, but analogous structures called tunneling nanotubes have been observed to facilitate intercellular communication between cancer cells (e.g. in pancreatic, glioblastoma, ovarian, and advanced lung cancer and mesothelioma (Cole et al., 2021; Desir et al., 2018; Matejka et al., 2024; Roehlecke and Schmidt, 2020), helping malignant cells thrive and survive in challenging environments.

### The role of KIF23 and ring canals in the evolution of cellular connectivity in multicellular animals

The role of KIF23 in incomplete cytokinesis and the formation of intercellular bridges underscores its potential significance in the evolution of multicellularity. By promoting the formation of midbodies large enough to be remodeled into ring canals, KIF23 possibly contributed to the emergence of cellular connectivity that enabled cooperative behaviors foundational to multicellular organization. KIF23, as part of the deeply conserved centralspindlin complex (Glotzer, 2024), is well-positioned to act as a molecular scaffold that supports the reorganization of midbodies. By interacting with key incomplete cytokinesis-specific proteins, KIF23 could facilitate the recruitment and assembly of additional structural and regulatory proteins necessary for enlarging midbodies into structures robust enough to be remodeled into ring canals. This process might involve cooperative interactions with actin-binding proteins, septins, or tubulin-associated complexes that stabilize the architecture of the midbody, ensuring its successful transition into a ring canal. Alternatively, post-translational modifications or unique regulatory mechanisms specific to incomplete cytokinesis could modulate KIF23’s activity, enabling its specialized role in the formation of syncytia.

The conserved enrichment of KIF23 protein in cells forming ring canals across taxa highlights its ancestral role in supporting syncytial systems and intercellular communication. The presence of KIF23-positive ring canals in interstitial stem cells and undifferentiated nematoblast clusters in *Hydra* exemplifies how such structures can enable the coordination of cell division and differentiation, critical processes in tissue organization. Similarly, in most choanoflagellates, the sister group to metazoans, intercellular bridges connect cells in rosette colonies (Laundon et al., 2019) and orthologs of KIF23 have been unambiguously identified in numerous Holozoan species (Glotzer, 2024). Furthermore, altered cytokinesis likely underlies intercellular bridge formation in choanoflagellates (Laundon et al., 2019) suggesting that the mechanisms for cellular connectivity and coordination were present even in unicellular ancestors. While the specific role of KIF23 has not been tested in any non-bilaterian or non-metazoan, its conserved expression in these syncytial systems suggests its likely contribution to the evolution of cellular networks that support multicellular complexity and the framework for cellular cooperation, synchrony, and specialization.

Further supporting this idea, our EdU incorporation assays in *Hydra* demonstrate that connected cells exhibit synchronous cell cycles, a feature likely mediated by intercellular bridges and one that may have conferred significant advantages in the evolution of multicellularity. By maintaining identical cell cycle stages, syncytial cells could achieve coordinated division and differentiation, minimizing competition for resources and optimizing the timing of cellular processes critical for tissue development. Such synchronization would enable cells within a multicellular structure to function collectively, facilitating the transition from loosely associated unicellular organisms to tightly integrated multicellular systems. EdU incorporation, combined with the identification of clusters of cells exhibiting synchronized cell cycles, could serve as a powerful approach for identifying cells connected by intercellular bridges, providing insights into the presence and functional significance of these structures across diverse tissues and species.

### The utility of Hydra as a model for ring canal biology

To date, there has been a significant, and almost exclusive, focus on ring canal formation in bilaterian organisms such as *Drosophila*, zebrafish, and mice. Note, in these animals and virtually all others studied, ring canals are present in the germline, suggesting that syncytial organization is important for maintaining germline pluripotency (Spradling, 2024). While these models have provided invaluable insights into the mechanisms of incomplete cytokinesis, their limitations - such as the absence of a robust cell culture system capable of sustaining incomplete cell division indefinitely - have hindered the ability to address detailed biochemical and mechanistic questions. *Hydra vulgaris* offers a unique opportunity to fill this gap, serving as a surrogate cell culture system where syncytial cells grow in abundance, comprising roughly 70% of all cells in an adult polyp, and cells can be disassociated and manipulated. Complementing these experimental capabilities, *Hydra* is supported by extensive genomic and transcriptomic resources as well as established transgenic tools. Integrating these resources with other interdisciplinary approaches provides a powerful opportunity to dissect the molecular and cellular mechanisms underpinning ring canal formation and function, while also exploring their evolutionary origins and adaptations.

## Methods

### *Hydra* strains and culturing conditions

*H. vulgaris* strains AEP (Martin et al., 1997) and cnnos1::eGFP (Hemmrich et al., 2012) were used in this study. Animals were fed *Artemia nauplii* 2-3 times a week and were cultured at 18°C according to standard procedures (Lenhoff and Brown, 1970). Experimental animals were starved for 24 hours.

### Immunofluorescence

Whole mount immunofluorescence was performed as previously described (Juliano et al., 2014), with slight modification. *Hydra* polyps were relaxed in 2% urethane in *Hydra* medium for 90 seconds and then fixed in 4% PFA in *Hydra* medium for 10 minutes. Following which, samples were washed with PBS and permeabilized with 0.5% Triton X-100 in PBS. Samples were incubated with blocking solution (1% BSA; 10% normal goat serum; 0.1% Triton X-100 in PBS) for one hour at room temperature. Primary antibodies were diluted in blocking solution and samples were incubated in the primary antibody solution overnight at 4°C. Samples were washed 3 X 10 minutes at room temperature in 1% BSA in PBS + 0.5% Tween-20 while gently rocking. Samples were incubated in secondary antibody diluted in blocking buffer and incubated at room temperature in the dark for one hour whilst rocking. Samples were then washed 3 X 10 minutes while gently rocking in PBS. During the second wash, DAPI (Invitrogen) was added to achieve a final concentration of 1 μg/μl to stain nuclei. Slides were mounted using Prolong Diamond Antifade Mountant (Invitrogen).

For labeling of syncytial cells from dissociated *Hydra* polyps, tissue was macerated in a solution of glycerin: acetic acid: water (1: 1: 13) at room temperature, as previously detailed (David, 1973). Briefly, *Hydra* were placed in maceration solution at a volume of 100 μl per polyp. Tissue was incubated for no more than three minutes and then the tube was gently agitated until the tissues dissociated to generate a suspension of single cells. Cells were then spotted onto gelatin-coated slides (Fisher Scientific) and dried for at least three hours before proceeding. Immunolabeling was performed using the same protocol as described for whole-mount labeling.

The following antibodies were used in this study: anti-HyKIF23 1:500 (Price et al., 2023); anti-Hywi 1:500 (Juliano et al., 2014); anti-phosphotyrosine 1:500 (clone 4G10, Sigma-Aldrich); alpha Tubulin-FITC 1:10 (sc-32293, Santa Cruz Biotechnology); Alexa Fluor 647 Phalloidin 1:400 (A22287, ThermoFisher); Alexa Fluor 488 goat anti-mouse IgG 1:500 (A11001); Alexa Fluor 488 goat anti-rabbit IgG 1:500 (A11008); Alexa Fluor 568 goat anti-mouse IgG 1:500 (A11004).

### EdU incorporation

*Hydra* polyps were incubated in 100 μM EdU (5-ethynyl-2’-deoxyuridine) in *Hydra* medium for 48 hours at 18°C. Animals were then fixed in 4% PFA in *Hydra* medium for 10 minutes at room temperature and EdU-labeled nuclei were detected with the Click-iT™ EdU Alexa Fluor 647 Imaging Kit (C10340, Invitrogen). Samples were subsequently processed for whole mount immunostaining as detailed above.

### Imaging acquisition, quantification, and statistical analysis

Whole mount and isolated cell preparations were imaged using a Zeiss LSM 980 confocal microscope and Plan-Apochromat 63X oil immersion objective (1.4 NA). Z-stacks were acquired at 0.2 μm step size. Raw image files were adjusted for brightness and contrast in Fiji/ImageJ. Ring canal diameter measurements were made by performing a line scan across the ring canals in the *en face* orientation; the distance between the two highest pixel intensities was determined from the XY coordinates. Generation of plots and statistical analyses was performed in either GraphPad Prism or R Studio. Statistical tests used are listed in the figures, figure legends, and text, when appropriate.

### Transmission Electron microscopy

Transmission electron microscopy was done according to standard protocols (Holstein et al., 2010). Wild-type polyps were relaxed in 2% urethane in *Hydra* medium and fixed with a mixture of glutaraldehyde and osmium tetroxide in phosphate buffer on ice for 2 hours. Samples were dehydrated in an increasing series of acetone and embedded into EMBed812 epoxy resin. The 80-nm ultrathin sections were cut with an ultracut UCT microtome (Leica) using a Diatome Diamond knife (Diatome). Sections were mounted on grids and stained with lead citrate and examined with Libra 120 energy filter transmission electron microscope (Zeiss). Images were acquired with a 2 X2k high speed camera and an ImageSP software (Tröndle).

### Bioinformatic analyses

Published single-cell RNA-seq datasets were obtained from Schnitzler et al. (2024), Li et al. (2022b), Steger et al. (2022), and Chari et al. (2021). Orthologous genes were identified using BLAST (Basic Local Alignment Search Tool). The sequences of the genes of interest from *Hydra vulgaris* were used as queries against the genome of the target species. The top hits with the highest alignment scores and appropriate e-values were considered orthologs. The identified orthologous proteins were aligned using the MUSCLE algorithm implemented in Geneious Prime (version 2024.0.4) to confirm sequence similarity and domain conservation.

UMAP projections of single-cell RNA sequencing data were obtained from the *Hydra* Single-Cell Browser (www.research.nhgri.nih.gov/HydraAEP/SingleCellBrowser/), which provides precomputed embeddings based on the dataset published by Cazet et al. (2023). The dataset includes cells profiled using Drop-seq to sequence the transcriptomes of 24,985 single *Hydra* cells, including stem cells to terminally differentiated cells, with preprocessing and clustering performed as described in the original publication. Clusters were identified based on annotations provided by the original study.

The gene expression levels of *kif23* homologs and other genes of interest were assessed across different cell types in each represented species. Dot plots were generated to visualize gene expression patterns using the ‘DotPlot’ function. All analyses were conducted in R Studio (version 2024.04.0+735) using the Seurat package (version 4.4.0) along with other supporting packages including dplyr (version 1.1.4), ggplot2 (version 3.5.1), and corrplot (version 0.92).

## Acknowledgements

We thank members of the Cooley, Sumigray (Yale), Juliano (UC Davis), and Dunn (Yale) laboratories for their invaluable input and stimulating conversations informing the work presented here. Additionally, we thank the *Hydra* community and participants of the Cnidofest Zoom Seminars for their critical feedback and for experimental suggestions. We are especially grateful to Celina Juliano for technical guidance, the anti-Hywi antibody, sharing of *Hydra* cloning reagents, providing us with all *Hydra* strains used in this study. We thank members of the Cooley laboratory for their careful reading of the manuscript. This work was supported by the National Institutes of Health (F32GM136029 to K.P. and R35GM141961 to L.C.).

## Contributions

K.L.P. and L.C. conceived the project. K.L.P. designed, performed, and analyzed experiments and wrote the manuscript. K.L.P. and D.M.T. performed experiments and subsequent quantifications and cultured animals. K.M.C.A. performed experiments and cultured animals. J.M. provided organized datasets for the analysis of single-cell RNA sequencing. W.S. and B.H. provided transmission electron micrographs of *Hydra* syncytia. All authors discussed results, edited, and commented on the manuscript at every stage.

## Supplemental Figures

**Supplemental Figure 1.**
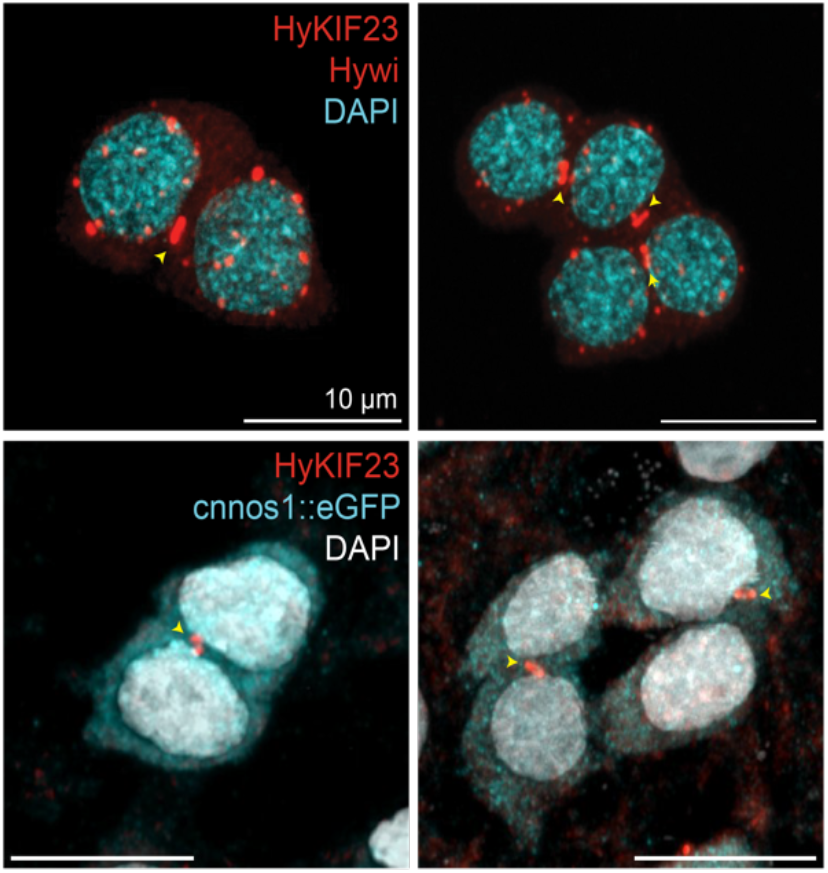
(Top) Interstitial stem cell pairs from macerated tissues were stained with anti-HyKIF23 to label ring canals (yellow arrowheads) and anti-Hywi to label perinuclear granules. (Bottom) Intact *Hydra* polyps expressing cnnos1::eGFP labeled with anti-HyKIF23.

**Supplemental Figure 2.**
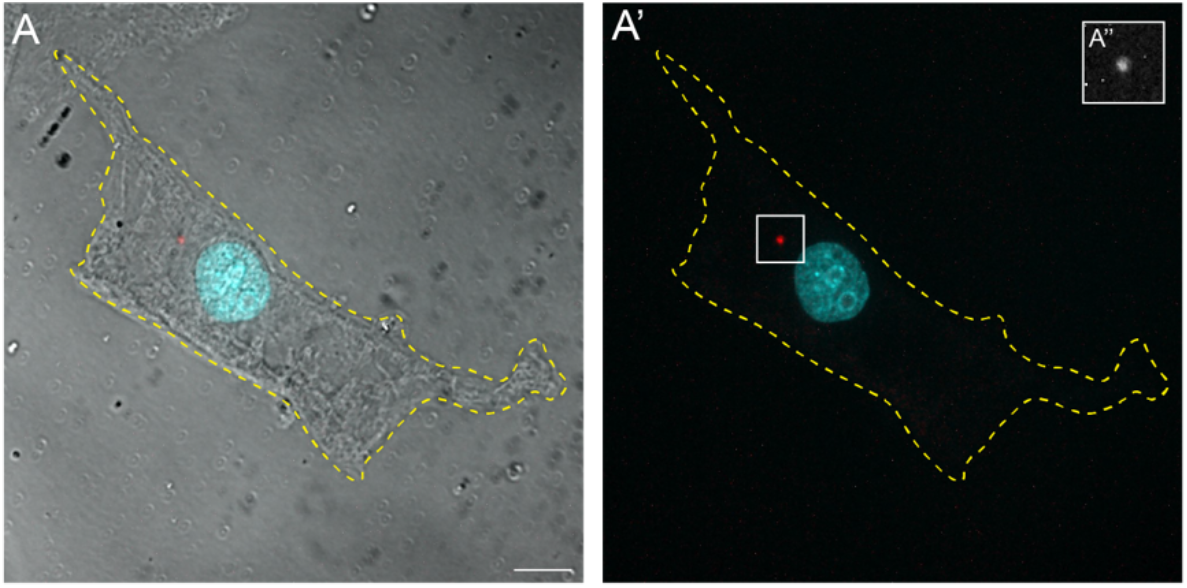
Brightfield (left) and immunofluorescence (right) images of a macerated epithelial cell stained with anti-HyKIF23 and DAPI. The HyKIF23-positive midbody (A’) remnant is present in the cytoplasm. Scale bar is 5 microns.

**Supplemental Figure 3.**
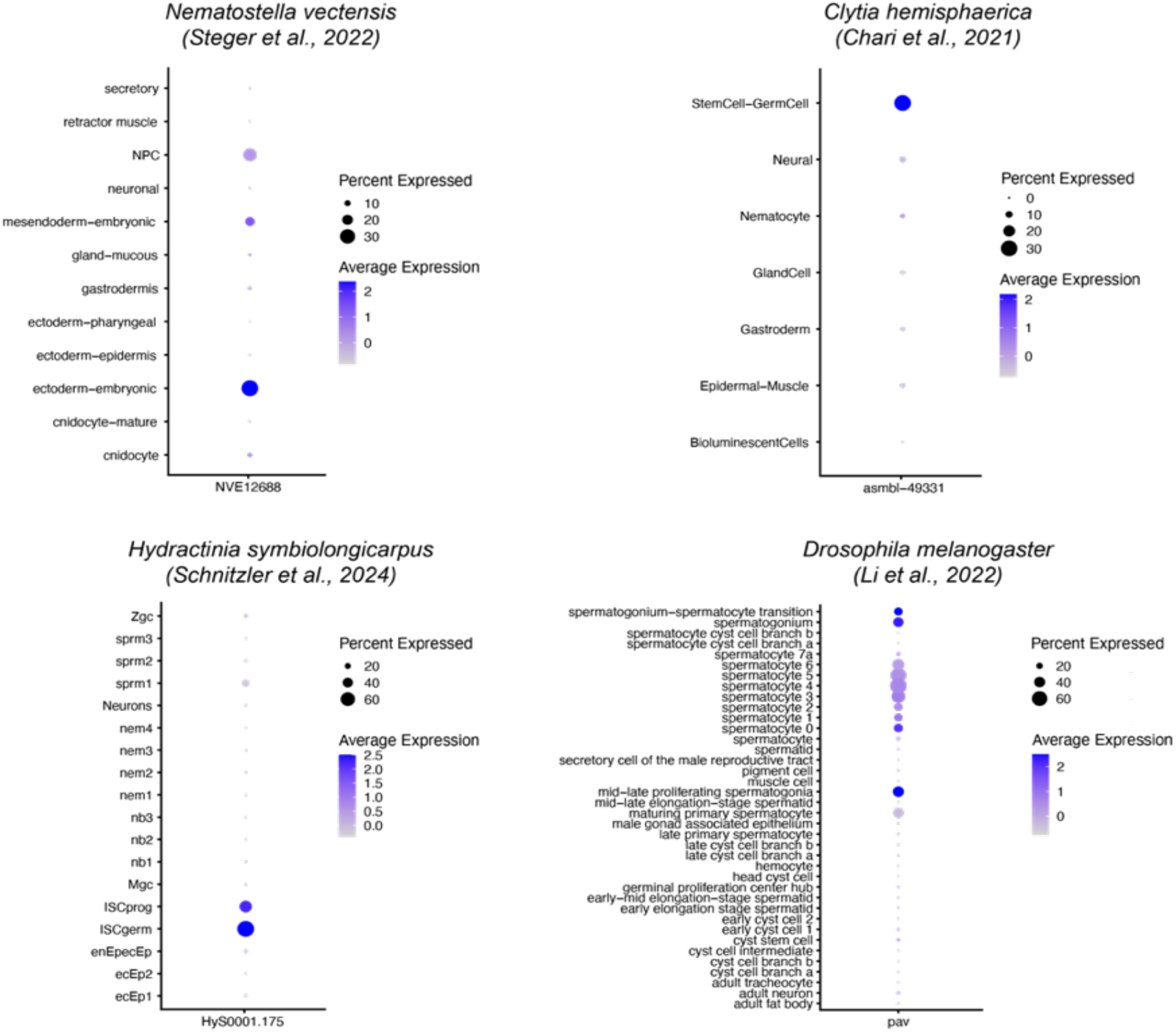
Expression of KIF23 homologs in other cnidarians as compared to Drosophila.

